# Evaluation of seasonal dynamics of fungal DNA assemblages in a flow-regulated stream in a restored forest using eDNA metabarcoding

**DOI:** 10.1101/2020.12.10.420661

**Authors:** Shunsuke Matsuoka, Yoriko Sugiyama, Yoshito Shimono, Masayuki Ushio, Hideyuki Doi

**Author notes:** Corresponding author: Shunsuke Matsuoka.

## Abstract

Investigation of seasonal variation in fungal communities is essential for understanding biodiversity and ecosystem functions. However, the conventional sampling method, with substrate removal and high spatial heterogeneity of community composition, makes surveying the seasonality of fungal communities challenging. Recently, water environmental DNA (eDNA) analysis has been explored for its utility in biodiversity surveys. In this study, we assessed whether the seasonality of fungal communities can be detected by monitoring eDNA in a forest stream. We conducted monthly water sampling in a forest stream over two years and used DNA metabarcoding to identify fungal eDNA. The stream water contained DNA from functionally diverse aquatic and terrestrial fungi, such as plant decomposers, parasites, and mutualists. The variation in the fungal assemblage showed a regular annual periodicity, meaning that the assemblages in a given season were similar, irrespective of the year or sampling. Furthermore, the strength of the annual periodicity varied among functional groups. Our results suggest that forest streams may act as a “trap” for terrestrial fungal DNA derived from different habitats, allowing the analysis of fungal DNA in stream water to provide information about the temporal variation in fungal communities in both the aquatic and the surrounding terrestrial ecosystems.

## Introduction

Fungi comprise diverse functional groups, such as decomposers, mutualists, and parasites, and thus play an important role in regulating ecosystem functions by driving the biogeochemical cycling of ecosystems and influencing the population dynamics of other organisms (Peay et al., 2016; Grossart et al., 2019). As the functional capacities of fungi may often differ among species (Nara, 2006; Osono, 2020; Zanne et al., 2020), it is essential to explore the spatiotemporal patterns of fungal communities, and the factors that shape them, to understand and predict how fungus-driven ecosystem functions respond to environmental changes (Lilleskov and Parrent, 2007). Previous studies have shown the spatial and temporal turnover of fungal communities in a variety of substrates, such as soils, living/dead plant tissues, and animal digestive tracts. For example, the species composition of fungi in soils varies at spatial scales of a few tens of centimeters to several meters (Bahram et al., 2016; Peay et al., 2016) or at time scales of several months (Koide et al., 2007; Pickles et al., 2010; Voříšková et al., 2013; Matsuoka et al., 2016; Sugiyama et al., 2020). In particular, the recent proliferation of high-throughput sequencing techniques has facilitated the simultaneous processing of a large number of samples in parallel, and information about the spatial variation of terrestrial and aquatic fungi in various ecosystems is rapidly accumulating (Peay et al., 2016; Grossart et al., 2019; Nilsson et al., 2019).

Typical factors causing temporal changes in the community include seasonal fluctuations in climate (e.g., temperature) and host phenology. Although studies on temporal changes in terrestrial and aquatic fungal communities are limited, several reports have demonstrated seasonal changes in community composition and the occurrence of fruiting bodies (Koide et al., 2007; Pickles et al., 2010; Sato et al., 2012; Voříšková et al., 2013; Taylor and Cunliffe, 2016). For example, saprotrophic fungi associated with tree leaves show seasonal patterns in response to the supply of tree foliage as a substrate (e.g., Voříšková et al., 2013). If the occurrence of fungal species and their associated fungal functions change seasonally, a one-time community survey (i.e., snapshot) cannot capture the fungal diversity and function at a study site and may lead to misestimations of diversity and function. For example, if the fungal community shows changes in response to seasonal defoliation events, the community composition may show a one-year periodicity. To detect such seasonal patterns, a continuous, multi-year monitoring study is needed (Bahram et al., 2014; Matsuoka et al., 2016; Sugiyama et al., 2020); however, as these surveys are limited, they hinder our understanding of the fungal community seasonality.

Investigating seasonal variation in fungal diversity based on time series sampling requires ingenuity in sampling, particularly in terrestrial habitats. Fungal communities in terrestrial habitats have been surveyed by collecting substrates from the field (Lindahl et al., 2013; Osono, 2014). Destructive sampling (i.e., a collected substrate is permanently lost from a study site) is a common method for investigating fungal community structure. Previous studies have shown that the fungal community composition in soils and plant substrates exhibits a high degree of spatial heterogeneity, where turnover can occur at scales of tens of centimeters to several meters (Bahram et al., 2016; Peay et al., 2016). Such compositional turnover is caused not only by environmental heterogeneity (e.g., soil chemistry and host plants) but also by stochastic factors, such as dispersal limitation and drift (Bahram et al., 2016; Peay et al., 2016). The large spatial turnover in community composition means that, to reflect the entire community, multiple samples must be taken from different places to represent the fungal community structure at the site. The collection and processing of many samples are not only expensive and time-and labor-consuming but can also cause disturbances in the form of substrate removal from the study area. Therefore, time series sampling may need to be designed to address these challenges.

Aquatic environmental DNA (eDNA) analysis may be used to overcome these difficulties. In recent years, the utility of aquatic eDNA surveys in biodiversity research has been explored, as this method allows for the detection of the DNA of organisms not only in water but also in surrounding terrestrial areas (Deiner et al., 2016; Khomich et al., 2017; LeBrun et al., 2018; Matsuoka et al., 2019). For example, Matsuoka et al. (2019) reported that the DNA of both aquatic and terrestrial fungi can be detected in river water in a forest landscape. This may be because the river water contains spores and mycelial fragments of fungi that have entered it from the surrounding terrestrial areas (Voronin, 2014). Furthermore, the fungal DNA assemblages are spatially structured; that is, similar DNA assemblages are found between rivers that are geographically close to each other and within a single tributary, but not between other tributaries (Matsuoka et al., 2019). This suggests that the investigation of fungal DNA assemblages, including those of both aquatic and terrestrial fungi, in river water may provide information on the diversity in the catchment area. Owing to these characteristics of aquatic eDNA, using these surveys in river and stream water may be potentially useful for addressing the above-mentioned challenges. First, water sampling does not involve the removal and destruction of terrestrial substrates such as soils; therefore, disturbance to an ecosystem is minor compared with that caused by conventional survey methods. Second, spatial heterogeneity is likely to be much lower than that in terrestrial substrates because of the high mobility of stream water. Indeed, the usefulness of aquatic eDNA in capturing seasonal community dynamics has been confirmed in aquatic invertebrates (Bista et al., 2017). However, it is also possible that seasonality of terrestrial fungal DNA may not be detectable in stream water because the concentration of terrestrial fungal DNA becomes lower due to dilution compared to terrestrial substrates such as in soil. However, the analysis of fungal DNA assemblages in water has only recently begun, and little is known regarding the seasonal patterns exhibited by the assemblages.

The objective of the present study was to investigate whether seasonal patterns of terrestrial fungi can be detected through time series sampling of forest stream water samples and fungal eDNA analysis. In particular, we addressed the following questions: (1) Which fungal DNA is detected in the forest stream water? (2) Do fungal DNA assemblages in water exhibit seasonality (e.g., annual periodicity)? (3) If the DNA assemblages show seasonal variation, is the variation related to climatic factors? This study was conducted in a flow-regulated stream within a restored forest fragment in an urban setting in Kyoto, Japan, with a particular focus on seasonal temporal variation. Here, the stream flow was controlled to maintain it within a certain range, depending on the amount of rainfall. In addition, unlike those in natural forest ecosystems, there were few large-scale disturbances, such as flooding. This setting allowed us to focus on seasonal changes, with minimal effects due to disturbances. Water sampling was conducted once a month for two years. Soil and plant tissues were also sampled at the study site to estimate the origin of fungal DNA detected in the stream water, although the sampling period did not overlap with that of water. The fungal assemblages in the water, soil, and plant samples were analyzed via DNA metabarcoding.

## Materials and methods

### Study site and sampling

The study site is a forest fragment located in Kyoto, in the western part of Japan (34.986751 ° N, 135.744950 ° E). It is a 30-year-old secondary growth forest (Fig. S1, approximately 0.6 ha) dominated by evergreen trees (Quercus glauca and Castanopsis sieboldii) and deciduous oak trees (Quercus serrata), which primarily comprise the natural vegetation in the area. A small stream runs through the forest. The main source of the stream water is the natural underground water of the Kyoto Basin. There is rich underground water flowing through the sand and gravel layer beneath the study area. This groundwater is pumped up from about 40 meters underground to provide a water source for the stream. The stream covers an area of 592 m2, and the daily water pumping volume is controlled to be approximately 43.5 m3, which translates to aboveground flow of approximately 0.5 L per second. Although it is difficult to determine the exact speed of water flow because the depth and width of the stream vary from place to place, the flow speed is gentle. Rainfall is another source of stream water. When rainfall is high, groundwater pumping is reduced to regulate flow. Stream water drains into the city sewer system from the lower reaches of the stream (near point D in Fig. S1) underground. The mean annual air temperature is 16.3 °C, and the mean annual precipitation is 1608.4 mm according to the Automatic Metrological Data Acquisition System (AMeDAS, Japan Meteorological Agency) at Kyoto Station, located 3 km northwest of the study site. This system automatically measures air temperature, precipitation, and the number of hours of sunshine on the ground and has been established at approximately 1,300 locations throughout Japan. The forest fragment is used as a biotope and is located more than 3 km away from the surrounding forests.

From December 2016 to November 2018, 1 L of surface water was collected using bleached plastic bottles once a month from each of the three sampling locations along the stream (The sampling locations are more than 40 meters away from each other, Fig. S1). The ammonium compound benzalkonium chloride (1 mL) was added in situ to prevent DNA degradation (Yamanaka et al., 2017). The bottles were stored in a cool, dark place and brought back to the laboratory for immediate filtration. The water samples were vacuum-filtered through 47 mm GF/F glass filters (pore size 0.7 μm, GE Healthcare, Little Chalfont, UK). The filters were stored at − 20 °C before DNA extraction. Total DNA was extracted from the filters using a PowerSoil Kit (Qiagen, Hilden, Germany). First, in a 2 mL tube, each filter was cut into small pieces using bleached dissecting scissors. Thereafter, the chopped filters were placed in bead tubes provided with the kit and vortexed for 20 min. DNA were then extracted according to the manufacturer’s instructions.

We calculated the mean daily air temperature (T), cumulative precipitation (P), and total hours of sunshine (S) for three days (3 d) and two weeks (2 w) prior to the survey date. Data on air temperature, rainfall, and total hours of sunshine during the study period were obtained from the AMeDAS Kyoto station (see above). The mean daily temperatures for 3 d and 2 w were expressed as T3d and T2w, respectively. These cumulative durations (3 d and 2 w) were selected based on previous studies and fungal life history (Matsuoka et al., 2016). We did not perform a comprehensive search of optimal cumulative durations to avoid high computational and runtime costs. The climatic variables at each sampling site are listed in Table S1.

In addition, to determine potential sources of fungal DNA detected in the stream water, we conducted additional soil and plant tissue (fresh and fallen leaves) sampling in the study forest in February 2021. First, one soil block of the fermentation-humus layer (10 cm × 10 cm each, approximately 5 cm depth), excluding the surface litter, was collected from each of the 10 sites. Three fresh and fallen leaves of two dominant evergreen species (Q. glauca and C. sieboldii) were collected from three sites. The collection sites are shown in Fig. S1. The collected samples were immediately brought back to the laboratory in cold bags for DNA extraction. In the laboratory, 250 mg of each soil sample was weighed out. The fresh leaves and fallen leaves were first rinsed and wiped with sterile water to remove particles adhering to the leaf surfaces. Then a 5 mm square piece of tissue was cut from each leaf and combined at each site. DNA was extracted from a total of 22 samples: 10 soil and 12 plant tissue samples (2 tree species × 2 types (fresh or fallen) × 3 sites). For comparison with the stream water samples, DNA was extracted using the same procedure as for the water samples using the PowerSoil Kit.

### Molecular identification of fungi

The procedures used for molecular experiments and bioinformatics were those described previously (Matsuoka et al., 2019). In brief, the fungal internal transcribed spacer 1 (ITS 1) region of rDNA was amplified using the ITS1-F-KYO2 (5’-TAG AGG AAG TAA AAG TCG TAA -3’) and ITS2-KYO2 (5’-TAG AGG AAG TAA AAG TCG TAA -3’) primer sets (Toju et al., 2012). The PCR amplicons were pooled and sequenced using the Illumina MiSeq platform at the Center for Ecological Research, Kyoto University, Japan. The sequence data were deposited in the Sequence Read Archive of the DNA Data Bank of Japan (accession number: DRA011182 and DRA012017). See Appendix 1 for details of the molecular analyses.

The raw MiSeq data were converted into FASTQ files using Bcl2gastq v2.18.0.12 and then demultiplexed using Claident pipeline (Tanabe and Toju, 2013; software available online: https://www.claident.org/). For all sequences, low quality ends with quality values below 30 were trimmed. For the forward and reverse sequences, those with overlaps were concatenated using the default settings of VSEARCH. If the sequence contained more than 10% of bases with qualities less than 30, the sequence was excluded. A total of 2,169,706 reads (30,135 ± 12,751 reads per sample, mean ± SE, n = 72) from water samples and 1,022,977 reads (46,499 ± 10,634 reads per sample, mean ± SE, n = 12) from soil and plant samples were assembled using Claident v0.2.2018.05.29. Potential chimeric sequences and sequencing errors were removed using UCHIME v4.2.40 (Edger et al., 2011) and algorithms in CD-HIT-OTU (Li et al., 2012), respectively. The remaining sequences were assembled at a threshold similarity of 97% (Osono, 2014), and the resulting consensus sequences represented the molecular operational taxonomic units (OTUs). For each of the obtained OTUs, taxonomic identification was conducted based on the query-centric auto-k-nearest-neighbor method (Tanabe and Toju, 2013) with the NCBI database, and subsequent taxonomic assignment was conducted using the lowest common ancestor algorithm (Huson et al., 2007) using Claident. The functional guild of each fungal OTU was estimated based on the FUNGuild database (Nguyen et al., 2016). One hundred OTUs (48,685 reads) that were identified as non-fungal organisms were discarded.

### Data analysis

The bioinformatic pipeline described above generated an OTU table (i.e., matrix of OTUs and samples with sequence reads in each cell entry). For this OTU matrix, cell entries with reads less than 0.0002% of the total reads in each sample (which corresponds to 2–13 reads, a typically used criterion in eDNA metabarcoding studies, Table S2) were removed because these rare entries could represent contamination. After this process, 1,935,576 reads and 4,409 OTUs were obtained. The number of sequences, taxa, functional groups, and consensus sequences of the obtained OTUs is shown in Table S2. All analyses were performed using R v.3.4.3 (R Core Team 2017). We confirmed that the number of sequence reads was enough to detect OTUs in the samples using the ‘rarefy’ and ‘rarecurve’ functions of the ‘vegan’ package (Fig. S2). Thus, we used raw data for further analyses without rarefying the data. The presence or absence of OTUs was recorded. Binary data were used for all the statistical analyses.

To test whether OTU compositions differed across sampling occasions and three sampling points, we conducted permutational multivariate analysis of variance (PERMANOVA), using the ‘adonis2’ command in the ‘vegan’ package ver. 2.5-6 with 9999 permutations. The presence/absence of the OTU data for each sample (n = 72) was converted into a dissimilarity matrix using the Raup-Crick index with 9999 permutations. The Raup-Crick dissimilarity index is calculated based on the null model approach, which is akin to the standardized effect size (SES) value described below. The community dissimilarity matrix was visualized via nonmetric multidimensional scaling (NMDS) using ‘ordinate’ and ‘plot_ordination’ commands in ‘phyloseq’ package ver. 1.28.0.

Subsequent analyses were performed for all OTUs and for each functional group (i.e., saprotrophic, mutualistic, and parasitic fungi). To evaluate temporal variations in OTU compositions among sampling occasions, the presence/absence data for each sampling occasion were merged. First, the effects of sampling year (first or second year) and month (i.e., twelve categories) on the OTU composition were tested with PERMANOVA (‘adonis2’ command, 9999 permutations). The presence/absence of the OTU data for each sampling occasion (n = 24) was converted into a Raup-Crick dissimilarity matrix with 9999 permutations. Next, the temporal dynamics of the OTU compositions were tested. The OTU matrix was converted into a dissimilarity matrix using the SES of the Jaccard dissimilarity index. The SES was defined as (Disobs–Disnull)/Dissd, where Disobs is the observed dissimilarity, Disnull is the mean of the null distribution of dissimilarity, and Dissd is the standard deviation of the null distribution. The null distribution was calculated based on 9999 randomizations, preserving both the site of occurrence and the OTU richness with ‘randomizeMatrix’ command in the ‘picante’ package. Temporal changes in SES values were tested by comparing SES values and temporal distances via generalized additive models (GAMs) using the ‘gam’ command in the ‘mgcv’ package ver. 1.8-31.

Finally, to estimate the contribution of climatic and temporal factors to the temporal changes in OTU composition, variation partitioning based on a distance-based redundancy analysis was conducted using the ‘capscale’ command in the ‘vegan’ package. The relative weight of each fraction (pure and shared fractions and unexplained fractions) was estimated using the methodology described by Peres-Neto et al. (2006). The Raup-Crick dissimilarity matrix was used for each sampling occasion. The detailed methods for variation partitioning are described in Matsuoka et al. (2016). Briefly, we constructed two models, including climatic and temporal distance variables, by applying the forward selection procedure (999 permutations with an alpha criterion = 0.05) of Blanchet et al. (2008). The full climatic model included six climatic variables (i.e., T3d + T2w + P3d + P2w + S3d + S2w). We then constructed a model using temporal distance vectors calculated using Moran’s eigenvector maps (MEM, Borcard et al. 2004). The MEM analysis produced a set of orthogonal variables derived from a temporal distance matrix, and MEM vectors represented various temporal patterns, such as periodicity. First, we created a temporal distance matrix between each sampling occasion based on the sampling dates. Next, MEM vectors were calculated from the temporal distance matrix using the ‘dbmem’ command in the ‘adespatial’ package. We used the 11 MEM vectors that best accounted for autocorrelation and then conducted forward selection (Table S1). Based on these two models, variation partitioning was performed by calculating the adjusted R2 values (Peres-Neto et al., 2006).

## Results and Discussion

### Fungal diversity in the stream water

Seventy-two water samples were obtained through a two-year (24 times × three sampling locations) survey during 2016–2018. A total of 1,935,576 reads from 72 samples were grouped into 4,409 operational taxonomic units (OTUs) with 97% sequence similarity (Table S2). The number of OTUs per sample was 350 ± 109 (mean ± standard error [SE]). Of these, 2553 OTUs (1,314,923 reads) were assigned as fungi. FUNGuild assigned 1,250 OTUs (805,598 reads. 28.4% of the total number of OTUs and 41.6% of the total reads) to functional guilds, 609 OTUs (355,218 reads) of which were saprotrophs; the others included parasites (261 OTUs, 147,417 reads) and mutualists (111 OTUs, 39,717 reads) (Table S2, Fig. S4). The remaining 269 OTUs included endophytes and OTUs with multiple functions. The major saprotrophs were plant saprotrophs (205 OTUs, 87,327 reads), including those considered to be aquatic hyphomycetes (Alatospora and Tetracladium) and terrestrial wood and leaf saprotrophs (e.g., Ganoderma and Mycena). The major parasites were plant pathogens (192 OTUs, 129,179 reads), such as Taphrina and Ciboria. Among the mutualists, the major fungi were ectomycorrhizal fungi (91 OTUs, 38,138 reads), such as Cortinarius and Russula. These plant parasites and ectomycorrhizal fungi are terrestrial fungi, validating our expectation that terrestrial eDNA could be detected in stream water. The detailed information on taxonomic and functional groups among the fungal OTUs are shown in Tables S2 and S3 and Figs. S3 and S4.

### Comparison of the fungal DNA detected in stream water and terrestrial substrates

Additional surveys of soil and vegetation to estimate the source of eDNA detected in the stream water were conducted in February 2021, and 974 and 394 of the fungal OTUs were detected in soil and vegetation, respectively (Table S2). Of the OTUs detected in water, 596 (13.5%) were detected in soil or plant materials, although the sampling period for soil and plant materials did not overlap with that of water. Of these, 352 OTUs were shared between water and soil, 184 were shared between water and plants, and 60 were shared between water, soil, and plants. For each water sampling occasion, about 20% of the OTUs and 10-70% of the reads detected in the water overlapped with those detected in the soil and plants (Fig. S5). The proportion of overlap increased in winter (from November to February). The percentages of taxonomic and functional groups among the fungal OTUs detected in the soil and plants are shown in Figs. S3 and S4.

Although the soil and plant tissue surveys were performed only once, the results suggested that DNA of terrestrial fungi present in the soil and plant tissues at the study site may be recruited into the stream. For these terrestrial fungi, spores and/or mycelial fragments released on land may have entered the stream through air, rainfall, and host tissues (Galante et al., 2011; Peay et al., 2012; Voronin, 2014; Chen et al., 2018; Castaño et al., 2019; Redondo et al., 2020).

### Temporal patterns of fungal OTU compositions and related variables

A total of 72 aquatic samples were analyzed, and the OTU composition differed significantly by sampling occasion (PERMANOVA, F = 1.368, R2 = 0.396, P = 0.0001). Although we sampled at three locations (corresponding to U, M, and D in Fig. 1) within the stream in each sampling occasion, the OTU composition did not differ significantly among sampling locations (PERMANOVA, F = 1.159, R2 = 0.033, P = 0.073). The result of NMDS ordination also showed temporal variation with the sampling occasion and month (Fig. 1). These results suggest that the DNA assemblages changed over time, and for a given sampling occasion, the differences in the DNA assemblages between sampling locations were smaller than the differences in time (i.e., spatial variation was less than temporal variation).

**Figure 1.**
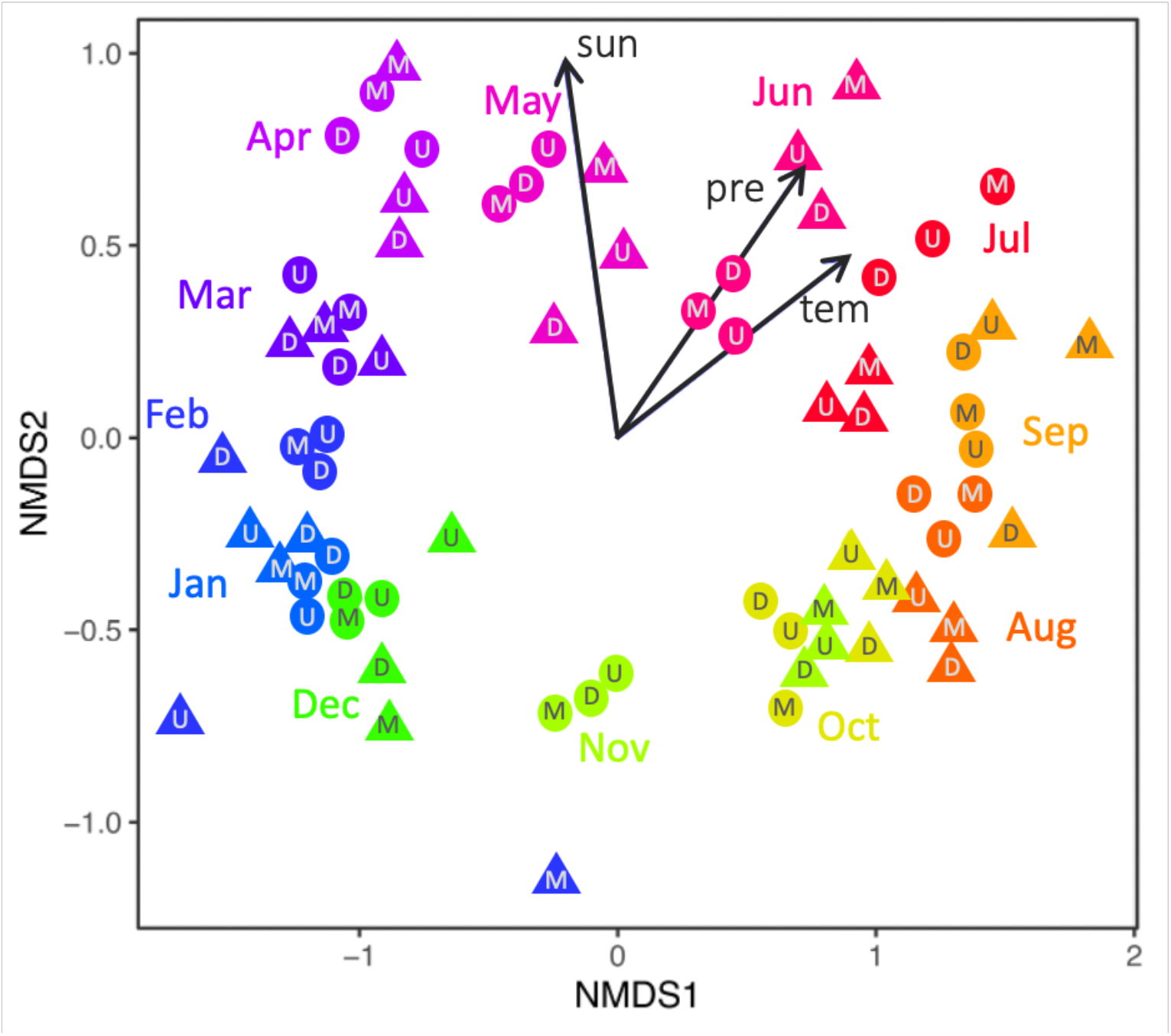
Dissimilarity in the DNA assemblages among sampling occasions, as revealed via nonmetric multidimensional scaling (NMDS) ordination (Raup-Crick index, stress value = 0.1412). Circles indicate the first year of sampling, and triangles indicate the second year of sampling. The colors indicate sampling months. The characters within the points indicate the collection location. The location indicated by each character is shown in Figure S1. Arrows denoted by “tem”, “pre”, and “sun” represent the effects of average air temperature, accumulated precipitation, and total hours of sunshine two weeks before each sampling date at the study site, respectively (see Table S1).

Subsequent analyses were performed on the presence/absence data for each sampling occasion (n = 24) by merging the presence/absence data in each sample for the same sampling occasion. The results of PERMANOVA showed that the OTU composition differed significantly by sampling month, but not by sampling year (Fig. 1 and Table S4; PERMANOVA, month, P = 0.0001, year, P = 0.461). A similar temporal pattern was observed in the results of the GAM regressions (Table S5), where the dissimilarity of OTU composition (SES value of Jaccard index) showed a one-year periodicity (Fig. 2). The local maximum values of community dissimilarity were observed at approximately 180 and 540 days, whereas the local minimum values were at 370 days. In addition, the dissimilarity, although periodic, tended to increase over time (as the x-axis increased). For example, the overall dissimilarity values were higher after approximately 600 days than after approximately 180 days. These results indicate the seasonal shifts in DNA assemblages were similar between years.

**Figure 2.**
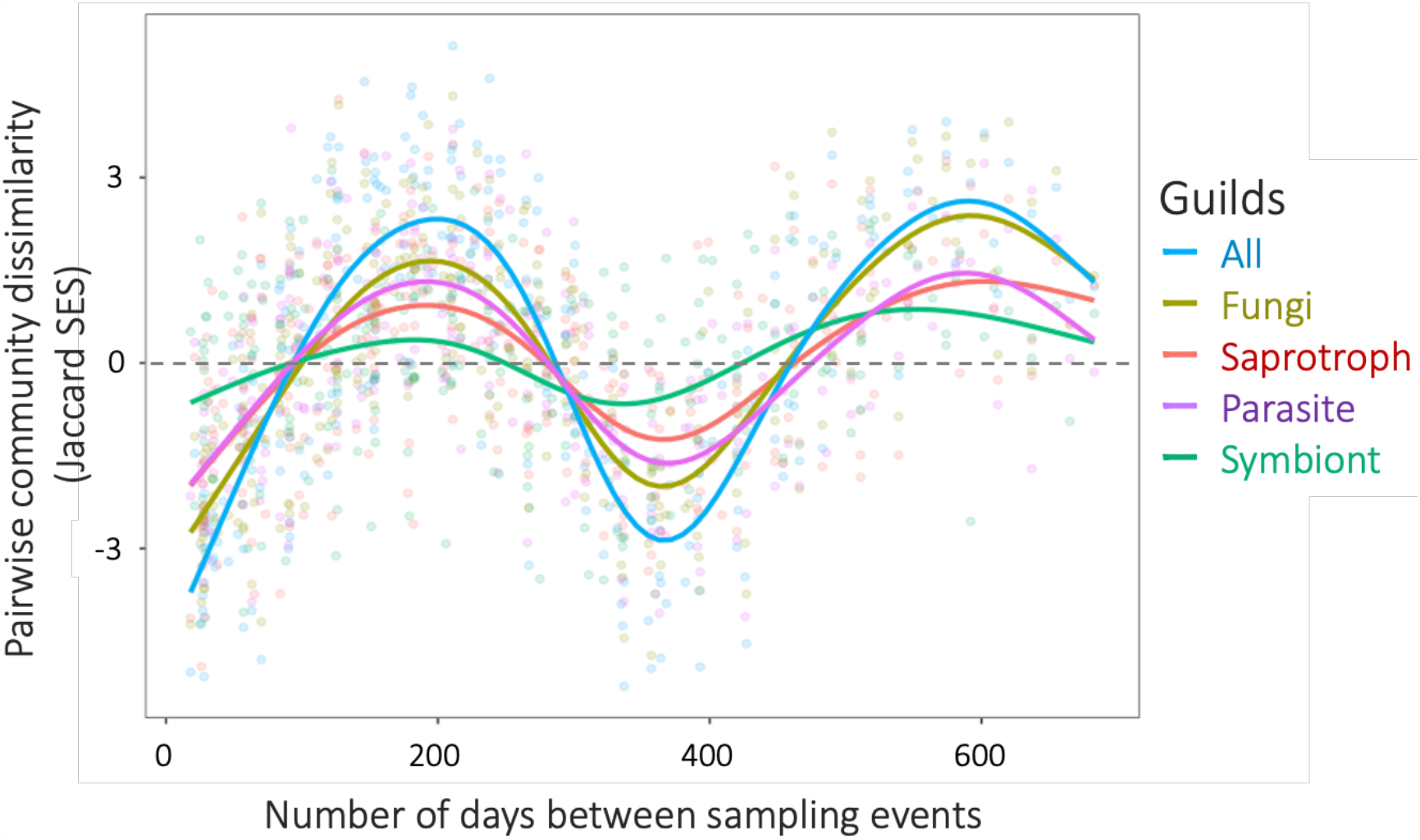
The relationship between the dissimilarity of OTU composition and temporal distance revealed via GAM. “All” shows the results for the dataset containing all detected OTUs, and “Fungi” shows the dataset containing only OTUs assigned as fungi. The regressions had significant coefficients for All, Fungi, and each functional group (P < 0.0001, see Table S5).

Temporal patterns of OTU composition showed a similar trend in individual guilds and the fungal OTU dataset. The OTU composition differed from month to month, regardless of the year in all guilds of fungi (Table S4, PERMANOVA, P < 0.05). Furthermore, the dissimilarity of OTU composition in all guilds showed a one-year periodicity (Fig. 2). The strength of periodicity varied across guilds; the explanatory power of sampling months in PERMANOVA, the periodicity of OTU composition dissimilarity (SES value), and the explanatory power of GAM were lower for mutualists than for saprotrophic and parasitic fungi (Tables S4 and S5, Fig. 2). The occurrence patterns of individual OTUs varied. First, there were OTUs that showed seasonal occurrence that were detected in the same month of different years. For example, OTU_0007 (Ramularia sp., plant pathogen) was detected from November to February both years, and OTU_2765 (Filosporella sp., saprotroph) was detected from February to May both years. However, many OTUs did not have a cyclic pattern of occurrence. For example, OTU_0646 (Mycena sp., saprotroph) and OTU_1103 (Inocybe sp., mutualists) were detected only in the first year. In addition, 2,171 OTUs (49%) were detected in only one sampling occasion over the two years. To provide an overview of the pattern of occurrence of individual OTUs, the typical occurrences of OTUs in each guild are shown in Fig. S6.

Variation partitioning was used to quantitatively estimate the relationships between climatic and temporal variables and the temporal changes in OTU composition (all fungal OTUs and individual guilds). For the climate variable, the mean daily air temperature for two weeks preceding each sampling date (T2w) was selected as a significant variable for all OTUs and for all guilds. In addition, the total number of hours of sunshine for the two weeks prior each sampling date (S2w) was selected for all OTUs, and cumulative precipitation for the three days prior each sampling date (P3d) was selected for the saprotrophic fungi. The relationship between OTU composition and the time was estimated using temporal distance vectors (Moran’s eigenvector maps, MEM), which represent various time patterns. For the MEM vectors, MEM3 and MEM4 were selected as significant vectors for all OTUs and for all guilds. These vectors represented an approximately one-year periodicity (Table S1). A list of selected variables is presented in Table S3. The climatic and temporal distance variables explained 6.6–16.8% and 6.3–24.7% of the OTU composition, respectively (Fig. S7). Of the fractions explained, 60–99% were shared between climate and temporal distance variables (Fig. S7).

These temporal patterns in the fungal DNA assemblages showed some agreement with known temporal patterns of terrestrial fungal community and/or spore release. First, the relatedness of temporal dynamics of community composition and climate could partly reflect the seasonality of fungal occurrence corresponding to the phenology of host organisms, including plants, plankton, and insects (e.g., Osono, 2008; Voříšková et al., 2013; Grossart et al., 2019). In the present study, a number of plant saprotrophic and parasitic fungi were detected (e.g., OTU_0007, Ramularia sp., this plant pathogen has been detected in both water and plant tissue). These fungal DNA can enter the stream along with the host tissue (e.g., tree leaves) (Voronin, 2014). In this case, temporal variations in fungal DNA in the stream may reflect plant phenology, such as seasonal increase in host species abundance and defoliation. These phenologies often correspond to climate seasonality, such as temperature variations (Cleland et al., 2007; Doi and Takahashi, 2008; Kitayama et al., 2021). In fact, tree defoliation increased at the study site from November to February, when air temperature and sunshine hours decreased. Second, spore release (i.e., formation of fruiting bodies) by fungi in the surrounding area could also be a source of seasonality. In a forest near the study site (4 km east), formations of fungal fruiting bodies have been reported to show seasonality, partly explained by seasonal changes in climate, including temperature (Sato et al., 2012). For example, in the area around the study site, the occurrence of fruiting bodies of Boletus species is concentrated during July–September (Sato et al., 2012); in fact, the DNA of several Boletus species has been detected in the stream water between July and September (e.g., OTU_1094 and 1136, Table S2). However, most of the temporal patterns of the DNA assemblages remain unexplained (Fig. S7). This may be partially because many fungal DNAs detected did not have distinct seasonality (Fig. S6); another factor could be the other environmental variables that affect the DNA assemblage, such as seasonal changes in water quality.

### Potential biases and limitations in the present approach

This study is based on simple monthly-monitoring in a flow-regulated stream, and many OTUs were not assigned to taxonomic or functional groups. Therefore, the generality of our findings, the factors that cause distinct periodicity in the community, and the contribution of individual taxa/functional groups to the temporal patterns of the community remain open questions. For example, if fungal DNA in the stream water reflects the fungal diversity of the surrounding terrestrial area and if the fungal DNA enters the stream mainly with rainfall, then the temporal pattern of fungal DNA in the stream could be influenced by the area surrounding the study site, the heterogeneity of the forest, and the seasonality of climate variables, including rainfall and wind. Also, the source and status of the fungal DNA detected in water (e.g., spores or mycelia), rates of DNA recruitment, movement (settling), and degradation in water, and the DNA concentrations in water remain to be determined. In addition, fungal spores can travel thousands of kilometers, although in small amounts (Wilkinson et al., 2012). Therefore, it is necessary to compare the spatial and temporal changes in fungal DNA assemblages in stream water with those of substrates other than the surrounding water.

While our results show the clear seasonality in fungal DNA assemblages in stream water, there may be some biases in our approach. In the present study, for example, the number of OTUs belonging to the phylum Glomeromycota was low. This could be related to the experimental procedure and the ecology of the fungus. First, there may be a taxa-specific detection bias due to match/mismatch of the PCR primers (Nilsson et al., 2019). Also, there are known examples of how experimental conditions (e.g., DNA extraction methods) and sequencing platforms affect metabarcoding results (Lindahl et al., 2013; Song et al., 2015; Ushio, 2019). Another reason may be that Glomeromycota do not produce airborne spores. Clarifying how experimental conditions and fungal ecology affect the detection of fungal DNA in water would contribute to more accurate monitoring of fungal community assemblages via stream water.

## Conclusion

In the present study, two-year monitoring of eDNA in a flow-regulated stream revealed that fungal DNA assemblages in water exhibited cyclical temporal variation. Differences in OTU composition due to differences in water sampling locations were small when the sampling time was the same. Furthermore, similar to the results of previous studies, the fungal DNA assemblages in water contained fungal DNA presumed to be aquatic and terrestrial. Our results suggest that forest streams act as a “trap” for terrestrial fungal DNA derived from different habitats, and that studies on stream water may provide information on the temporal variation of fungal communities living not only in the water but also in the surrounding ecosystem. Despite several technical limitations, this method may enable long-term monitoring of fungal community compositions in forest ecosystems at lower sampling costs; moreover, disturbance to the system is minimal compared to that caused by conventional approaches that target individual substrates. An important future research topic will be the elucidation of dynamic processes, such as the recruitment into, and loss of fungal DNA from, water, as well as the quantification of temporal changes in individual OTUs, and the identification of causal factors (Ushio, 2019; Ushio, 2020).

## Acknowledgments

We thank: Yukihiro Morimoto, Keizo Tabata, the staff of the Kyoto Greenery Association, members of the monitoring group of Inotino Mori, and Chisako Sugiyama for their assistance with the field work; Hirotoshi Sato and Mariko Nagano for assistance in laboratory work; and Hirokazu Toju for help with the MiSeq sequence. This study received partial financial support from the Japan Society for the Promotion of Science (JSPS) to SM (20J01732). This study was supported by the Environment Research and Technology Development Fund (JPMEERF20164002) and a Joint Usage/Research of Center for Ecological Research, Kyoto University.

## Data Accessibility

The sequence data were deposited in the Sequence Read Archive of the DNA Data Bank of Japan (accession number: DRA011182).

## Author Contributions

SM and HD designed the study, and SM, YShimono, and YSugiyama conducted field sampling. SM, YSugiyama, and MU contributed to the molecular experiments. SM, YSugiyama HD, and MU analyzed the data and interpreted the results. SM wrote the initial draft of the manuscript. All authors critically reviewed the manuscript.

## Supporting information

**Table S1** Climate variables and temporal distance vectors at each sampling occasion.

**Table S2** List of all OTUs, the number of their sequence reads and consensus sequences, and taxonomic and functional assignments.

**Table S3** The summary of taxonomic assignments.

**Table S4** PERMANOVA results for the composition of DNA assemblages.

**Table S5** GAM results for the relationship between the compositional dissimilarity of OTU composition and temporal distance.

**Table S6** List of selected variables in variation partitioning.

**Figure S1**. A simple map (a) and a photograph (b) of the study site. (a) The blue line and red arrows indicate the stream and water sampling points, respectively. The stream is labelled upstream to downstream in the order U, M, and D. One grid indicates 10 m. S1– S10 indicate the soil sampling locations. Plant tissues were collected at three locations: S1, S3, and S10.

**Figure S2**. Relationship between the number of sequence reads and OTU numbers, i.e., rarefaction curves for the samples.

**Figure S3**. Phylum level proportions of the DNA assemblages of each sample. (a) Percentage of the number of OTUs, and (b) percentage of the number of reads in the sequence. The order of the samples is the same as that in Table S2.

**Figure S4**. Functional composition of the DNA assemblages of each sample. (a) Percentage of the number of OTUs, and (b) percentage of the number of reads in the sequence. The order of the samples is the same as that in Table S2.

**Figure S5**. Percentage of OTUs and reads detected in stream water that overlapped with OTUs and reads detected in (a) soils and (b) plant leaves. The results are shown for the dataset of all OTUs and the dataset of dominant OTUs (top 100 OTUs with high detection frequency) detected in the water.

**Figure S6**. Heatmaps showing the temporal occurrence patterns of the top 50 OTUs, with the number of sample occurrences for each guild. The OTUs were clustered using the word method based on their occurrence patterns. (a) Saprotroph, (b) Parasite, and (c) Mutualist.

**Figure S7**. Bar plots showing pure and shared effects of climatic and temporal variables on the fungal OTU assemblages, as derived from variation partitioning analysis. Numbers indicate the proportions of explained variations. “All” shows the results for the dataset containing all detected OTUs, and “Fungi” shows the results for the dataset containing only OTUs assigned as fungi.

## Appendix 1 The detailed procedure of molecular analyses

For MiSeq sequencing, the fungal internal transcribed spacer 1 (ITS 1) region of rDNA was amplified. The first-round PCR (first PCR) amplified the ITS1 region using the ITS1-F-KYO2 and ITS2-KYO2 primer set. An Illumina sequencing primer and six random bases (N) were combined to produce each primer. Thus, the forward primer sequence was: 5′-*ACA CTC TTT CCC TAC ACG ACG CTC TTC CGA TCT* NNNNNN TAG AGG AAG TAA AAG TCG TAA -3′ and the reverse primer sequence was: 5′-*GTG ACT GGA GTT CAG ACG TGT GCT CTT CCG ATC T* NNNNNN TTY RCT RCG TTC TTC ATC-3′. The italic and normal letters represent MiSeq sequencing primers and fungi-specific primers, respectively. The six random bases (N) were used to enhance cluster separation on the flowcells during initial base call calibrations on MiSeq. The 1^st^ PCR was performed in a 12 μl volume with the buffer system of KODFX NEO (TOYOBO, Osaka, Japan), which contained 2.0 μl of template DNA, 0.2 μl of KOD FX NEO, 6.0 μl of 2× buffer, 2.4 μl of dNTP, and 0.7 μl each of the two primers (5 μM). The PCR conditions were as follows; an initial incubation for 2 min at 94°C followed by 5 cycles of 10 s at 98°C, 30 s at 68°C for annealing and 30 s at 68°C, 5 cycles of 10 s at 98°C, 30 s at 65°C and 30 s at 68°C; 5 cycles of 10 s at 98°C, 30 s at 62°C and 30 s at 68°C; 25 cycles of 10 s at 98°C, 30 s at 59°C and 30 s at 68°C, and a final extension of 5 min at 68°C. Eight replicate first-PCRs (per sample) were performed to mitigate the reaction-level PCR bias. Then, the duplicated first PCR amplicons (per sample) were combined, resulting in a template per sample for the second PCR. The PCR templates were purified using Agencourt AMPure XP (PCR product: AMPure XP beads = 1:0.8; Beckman Coulter, Brea, California, USA) before the second PCR.

The second PCR amplified the first PCR amplicons using the primers (forward) 5′-*AAT GAT ACG GCG ACC ACC GAG ATC TAC AC* XXXXXXXX TCG TCG GCA GCG TCA GAT GTG TAT AAG AGA CAG-3′ and (reverse) 5′-*CAA GCA GAA GAC GGC ATA CGA GAT* XXXXXXXX GTC TCG TGG GCT CGG AGA TGT GTA TAA GAG ACA G-3′. The italic and normal letters represent the MiSeqP5/P7 adapter and sequencing primers, respectively. The 8X bases represent dual-index sequences inserted to identify different samples. The second PCR was carried out with 12 μl reaction volume containing 1.0 μl of template, 6 μl of 2× KAPA HiFi HotStart ReadyMix (KAPA Biosystems, Wilmington, Washington, USA), 1.4 μl of each primer (2.5 μM), and 2.2 μl of sterilized distilled water. The PCR conditions were as follows; an initial incubation for 3 min at 95°C followed by 12 cycles of 20 s at 98°C, 15 s at 72°C for annealing and extension, and a final extension of 5 min at 72°C.

The indexed second PCR amplicons were pooled to make a library to be sequenced on MiSeq. The volume of each sample added to the library was adjusted to normalize the concentrations of each second PCR product. The pooled library was purified using Agencourt AMPure XP. A target-sized DNA of the purified library (approximately 380–510 base pairs [bp]) was then excised using E-Gel SizeSelect (ThermoFisher Scientific, Waltham, MA, USA). The double-stranded DNA concentration of the library was then adjusted to 4 nmol/L using Milli-Q water, and the DNA sample was applied to the Illumina MiSeq platform at Kyoto University, Japan.

## References

Bahram, M., Kohout, P., Anslan, S., Harend, H., Abarenkov, K., & Tedersoo, L. (2016). Stochastic distribution of small soil eukaryotes resulting from high dispersal and drift in a local environment. The ISME Journal, 10(4), 885–896. https://doi.org/10.1038/ismej.2015.164

Bahram, M., Peay, K. G., & Tedersoo, L. (2014). Local-scale biogeography and spatiotemporal variability in communities of mycorrhizal fungi. New Phytologist, 205(4), 1454–1463. https://doi.org/10.1111/nph.13206

Bista, I., Carvalho, G., Walsh, K., Seymour, M., Hajibabaei, M., Lallias, D., Christmas, M. & Creer, S. (2017). Annual time-series analysis of aqueous eDNA reveals ecologically relevant dynamics of lake ecosystem biodiversity. Nature Communications, 8, 14087. https://doi.org/10.1038/ncomms14087

Blanchet, F. G., Legendre, P., & Borcard, D. (2008). Forward selection of explanatory variables. Ecology, 89(9), 2623–2632. https://doi.org/10.1890/07-0986.1

Borcard, D., Legendre, P., Avois-Jacquet, C., & Tuomisto, H. (2004). Dissecting the spatial structure of ecological data at multiple scales. Ecology, 85(7), 1826–1832. https://doi.org/10.1890/03-3111

Castaño, C., Bonet, J. A., Oliva, J., Farré, G., Martínez de Aragón, J., Parladé, J., Pera, J., & Alday, J. G. (2019). Rainfall homogenizes while fruiting increases diversity of spore deposition in Mediterranean conditions. Fungal Ecology, 41, 279–288. https://doi.org/10.1016/j.funeco.2019.07.007

Chen, W., Hambleton, S., Seifert, K. A., Carisse, O., Diarra, M. S., Peters, R. D., Lowe, C., Chapados, J. T., & Lévesque, C. A. (2018). Assessing performance of spore samplers in monitoring aeromycobiota and fungal plant pathogen diversity in Canada. Applied and Environmental Microbiology, 84(9), e02601–17. https://doi.org/10.1128/AEM.02601-17

Cleland, E. E., Chuine, I., Menzel, A., Mooney, H. A., & Schwartz, M. D. (2007). Shifting plant phenology in response to global change. Trends in Ecology & Evolution, 22(7):357–65. https://doi.org/10.1016/j.tree.2007.04.003.

Deiner, K., Fronhofer, E. A., Mächler, E., Walser, J.-C., & Altermatt, F. (2016). Environmental DNA reveals that rivers are conveyer belts of biodiversity information. Nature Communications, 7(1), 12544. https://doi.org/10.1038/ncomms12544

Doi, H. & Takahashi, M. (2008). Latitudinal patterns in the phonological responses of leaf colouring and leaf fall to climate changes in Japan. Global Ecology and Biogeography, 17, 556–561.

Edgar, R. C., Haas, B. J., Clemente, J. C., Quince, C., & Knight, R. (2011). UCHIME improves sensitivity and speed of chimera detection. Bioinformatics, 27(16), 2194–2200. https://doi.org/10.1093/bioinformatics/btr381

Galante, T. E., Horton, T. R., & Swaney, D. P. (2011). 95 % of basidiospores fall within 1 m of the cap: A field-and modeling-based study. Mycologia, 103(6), 1175–1183. https://doi.org/10.3852/10-388

Grossart, H.-P., Van den Wyngaert, S., Kagami, M., Wurzbacher, C., Cunliffe, M., & Rojas-Jimenez, K. (2019). Fungi in aquatic ecosystems. Nature Reviews Microbiology, 17(6), 339–354. https://doi.org/10.1038/s41579-019-0175-8

Huson, D. H., Auch, A. F., Qi, J., & Schuster, S. C. (2007). MEGAN analysis of metagenomic data. Genome Research, 17(3), 377–386. https://doi.org/10.1101/gr.5969107

Khomich, M., Cox, F., Andrew, C. J., Andersen, T., Kauserud, H., & Davey, M. L. (2018). Coming up short: Identifying substrate and geographic biases in fungal sequence databases. Fungal Ecology, 36, 75–80. https://doi.org/10.1016/j.funeco.2018.08.002

Khomich, M., Davey, M.L., Kauserud, H., Rasconi, S., & Andersen, T. (2017) Fungal communities in Scandinavian lakes along a longitudinal gradient. Fungal Ecology 27, 36–46. https://doi.org/10.1016/j.funeco.2017.01.008

Kitayama, K., Ushio, M., & Aiba, S-I. (2020) Temperature is a dominant driver of distinct annual seasonality of leaf litter production of equatorial tropical rain forests. Journal of Ecology, 109: 727–736. https://doi.org/10.1111/1365-2745.13500

LeBrun, E. S., Taylor, D. L., King, R. S., Back, J. A., & Kang, S. (2018). Rivers may constitute an overlooked avenue of dispersal for terrestrial fungi. Fungal Ecology, 32, 72–79. https://doi.org/10.1016/j.funeco.2017.12.003

Li, W., Fu, L., Niu, B., Wu, S., & Wooley, J. (2012). Ultrafast clustering algorithms for metagenomic sequence analysis. Briefings in Bioinformatics, 13(6), 656–668. https://doi.org/10.1093/bib/bbs035

Lilleskov, E. A., & Parrent, J. L. (2007). Can we develop general predictive models of mycorrhizal fungal community-environment relationships? New Phytologist, 174(2), 250–256. https://doi.org/10.1111/j.1469-8137.2007.02023.x

Lindahl, B. D., Nilsson, R. H., Tedersoo, L., Abarenkov, K., Carlsen, T., Kjøller, R., Kõljalg, U., Pennanen, T., Rosendahl, S., Stenlid, J., & Kauserud, H. (2013). Fungal community analysis by high-throughput sequencing of amplified markers-a user’s guide. New Phytologist, 199(1), 288–299. https://doi.org/10.1111/nph.12243

Matsuoka, S., Kawaguchi, E., & Osono, T. (2016). Temporal distance decay of similarity of ectomycorrhizal fungal community composition in a subtropical evergreen forest in Japan. FEMS Microbiology Ecology, 92(5), fiw061. https://doi.org/10.1093/femsec/fiw061

Matsuoka, S., Sugiyama, Y., Sato, H., Katano, I., Harada, K., & Doi, H. (2019). Spatial structure of fungal DNA assemblages revealed with eDNA metabarcoding in a forest river network in western Japan. Metabarcoding and Metagenomics, 3, e36335. https://doi.org/10.3897/mbmg.3.36335

Nara K. (2006) Ectomycorrhizal networks and seedling establishment during early primary succession. New Phytologist, 169(1), 169–78. https://doi.org/10.1111/j.1469-8137.2005.01545.x.

Nguyen, N. H., Song, Z., Bates, S. T., Branco, S., Tedersoo, L., Menke, J., Schilling, J. S., & Kennedy, P. G. (2016). FUNGuild: An open annotation tool for parsing fungal community datasets by ecological guild. Fungal Ecology, 20, 241–248. https://doi.org/10.1016/j.funeco.2015.06.006

Nilsson, R. H., Anslan, S., Bahram, M., Wurzbacher, C., Baldrian, P., & Tedersoo, L. (2019). Mycobiome diversity: High-throughput sequencing and identification of fungi. Nature Reviews Microbiology, 17(2), 95–109. https://doi.org/10.1038/s41579-018-0116-y

Osono, T. (2008). Endophytic and epiphytic phyllosphere fungi of Camellia japonica: seasonal and leaf age-dependent variations. Mycologia, 00(3), 387–391. https://doi.org/10.3852/07-110R1

Osono, T. (2014). Metagenomic approach yields insights into fungal diversity and functioning. In T. Sota, H. Kagata, Y. Ando, S. Utsumi, & T. Osono (Eds). Species Diversity and Community Structure (pp. 1–23). Springer.

Osono, T. (2020). Functional diversity of ligninolytic fungi associated with leaf litter decomposition. Ecological Research, 35(1), 30–43. https://doi.org/10.1111/1440-1703.12063

Peay, K. G., Kennedy, P. G., & Talbot, J. M. (2016). Dimensions of biodiversity in the Earth mycobiome. Nature Reviews Microbiology, 14(7), 434–447. https://doi.org/10.1038/nrmicro.2016.59

Peay, K. G., Schubert, M. G., Nguyen, N. H., & Bruns, T. D. (2012). Measuring ectomycorrhizal fungal dispersal: Macroecological patterns driven by microscopic propagules: measuring mycorrhizal fungal dispersal. Molecular Ecology, 21(16), 4122–4136. https://doi.org/10.1111/j.1365-294X.2012.05666.x

Peres-Neto, P. R., Legendre, P., Dray, S., & Borcard, D. (2006). Variation partitioning of species data matrices: estimation and comparison of fractions. Ecology, 87(10), 2614–2625. https://doi.org/10.1890/0012-9658(2006)87[2614:VPOSDM]2.0.CO;2

R Core Team (2017). R: A language and environment for statistical computing. Retrieved from https://www.R-project.org/

Redondo, M. A., Berlin, A., Boberg, J., & Oliva, J. (2020). Vegetation type determines spore deposition within a forest–agricultural mosaic landscape. FEMS Microbiology Ecology, 96(6), fiaa082. https://doi.org/10.1093/femsec/fiaa082

Sato, H., Morimoto, S., & Hattori, T. (2012). A thirty-year survey reveals that ecosystem function of fungi predicts phenology of mushroom fruiting. PLoS ONE, 7(11), e49777. https://doi.org/10.1371/journal.pone.0049777

Schoch, C. L., Seifert, K. A., Huhndorf, S., Robert, V., Spouge, J. L., Levesque, C. A., Chen, W., & Fungal Barcoding Consortium (2012). Nuclear ribosomal internal transcribed spacer (ITS) region as a universal DNA barcode marker for Fungi. Proceedings of the National Academy of Sciences, 109(16), 6241–6246. https://doi.org/10.1073/pnas.1117018109

Song, Z., Schlatter, D., Kennedy, P., Kinkel, L. L., Kistler, H. C., Nguyen, N., & Bates, S. T. (2015). Effort versus reward: preparing samples for fungal community characterization in high-throughput sequencing surveys of soils. PLOS ONE, 10(5), e0127234. https://doi.org/10.1371/journal.pone.0127234

Sugiyama, Y., Matsuoka, S., & Osono, T. (2020). Two-years of investigation revealed the inconsistency of seasonal dynamics of an ectomycorrhizal fungal community in Japanese cool-temperate forest across years. FEMS Microbiology Ecology, 96(7), fiaa118. https://doi.org/10.1093/femsec/fiaa118

Tanabe, A. S., & Toju, H. (2013). Two new computational methods for universal DNA barcoding: a benchmark using barcode sequences of Bacteria, Archaea, Animals, Fungi, and Land Plants. PLoS ONE, 8(10), e76910. https://doi.org/10.1371/journal.pone.0076910

Taylor, J. D., & Cunliffe, M. (2016). Multi-year assessment of coastal planktonic fungi reveals environmental drivers of diversity and abundance. The ISME Journal, 10(9), 2118–2128. https://doi.org/10.1038/ismej.2016.24

Tedersoo, L., Anslan, S., Bahram, M., Kõljalg, U., & Abarenkov, K. (2020). Identifying the ‘unidentified’ fungi: a global-scale long-read third-generation sequencing approach. Fungal Diversity, 103, 273– 293. https://doi.org/10.1007/s13225-020-00456-4

Toju, H., Tanabe, A. S., Yamamoto, S., & Sato, H. (2012) High-coverage ITS primers for the DNA-based identification of ascomycetes and basidiomycetes in environmental samples. PLoS ONE, 7, e40863. https://doi.org/10.1371/journal.pone.0040863

Ushio, M. (2019). Use of a filter cartridge combined with intra-cartridge bead-beating improves detection of microbial DNA from water samples. Methods in Ecology and Evolution, 10(8), 1142–1156. https://doi.org/10.1111/2041-210X.13204

Ushio, M. (2020). Interaction capacity underpins community diversity. bioRxiv. https://doi.org/10.1101/2020.04.08.032524

Voříšková, J., Brabcová, V., Cajthaml, T., & Baldrian, P. (2013). Seasonal dynamics of fungal communities in a temperate oak forest soil. New Phytologist, 201(1), 269–278. https://doi.org/10.1111/nph.12481

Voronin, L. V. (2014). Terrigenous micromycetes in freshwater ecosystems (review). Inland Water Biology, 7(4), 352–356. https://doi.org/10.1134/S1995082914040191

Wilkinson, D. M., Koumoutsaris, S., Mitchell, E. A. D., & Bey, I. (2012). Modelling the effect of size on the aerial dispersal of microorganisms: Modelling the aerial dispersal of microorganisms. Journal of Biogeography, 39(1), 89–97. https://doi.org/10.1111/j.1365-2699.2011.02569.x

Yamanaka, H., Minamoto, T., Matsuura, J., Sakurai, S., Tsuji, S., Motozawa, H., Hongo, M., Sogo, Y., Kakimi, N., Teramura, I., Sugita, M., Baba, M., & Kondo, A. (2017). A simple method for preserving environmental DNA in water samples at ambient temperature by addition of cationic surfactant. Limnology, 18(2), 233–241. https://doi.org/10.1007/s10201-016-0508-5

Zanne, A. E., Abarenkov, K., Afkhami, M. E., Aguilar-Trigueros, C. A., Bates, S., Bhatnagar, J. M., Busby, P. E., Christian, N., Cornwell, W. K., Crowther, T.W., Flores-Moreno, H., Floudas, D., Gazis, R., Hibbett, D., Kennedy, P., Lindner, D. L., Maynard, D. S., Milo, A. M., Nilsson, R. H., Powell, J., Schildhauer, M., Schilling, J., & Treseder, K. K. (2020) Fungal functional ecology: A trait-based approach to plant-associated fungi. Biological reviews of the Cambridge Philosophical Society, 95(2), 409–433. https://doi.org/10.1111/brv.12570.

