## Supplemental Figures for "Evaluation of seasonal dynamics of fungal DNA assemblages in a flow-regulated stream in a restored forest using eDNA metabarcoding"

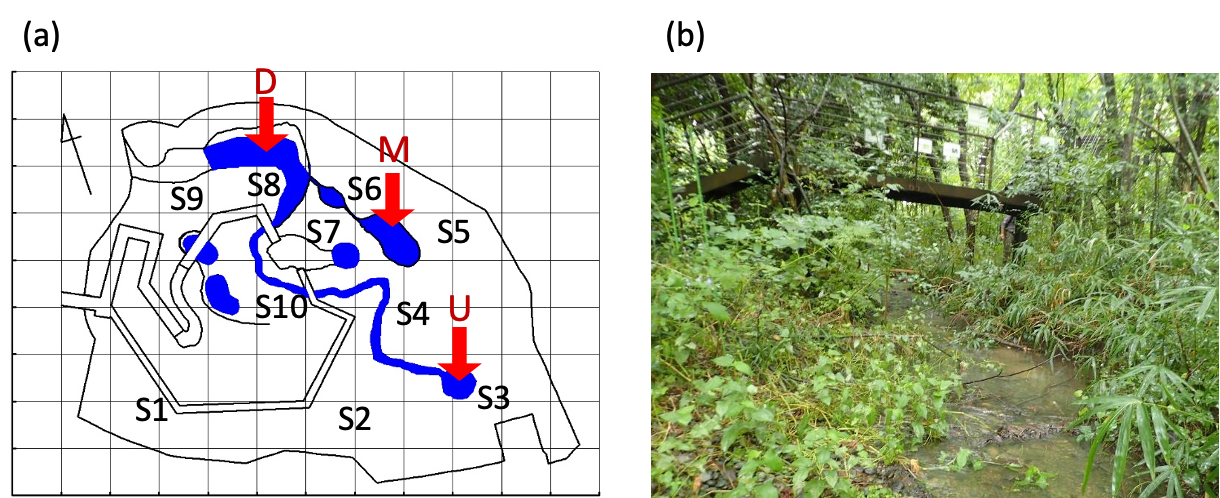

**Figure S1.** A simple map (a) and a photograph (b) of the study site. (a) The blue line and red arrows indicate the stream and water sampling points, respectively. The stream is labelled upstream to downstream in the order U, M, and D. One grid indicates 10 m. S1–S10 indicate the soil sampling locations. Plant tissues were collected at three locations: S1, S3, and S10.

(Figure S2)

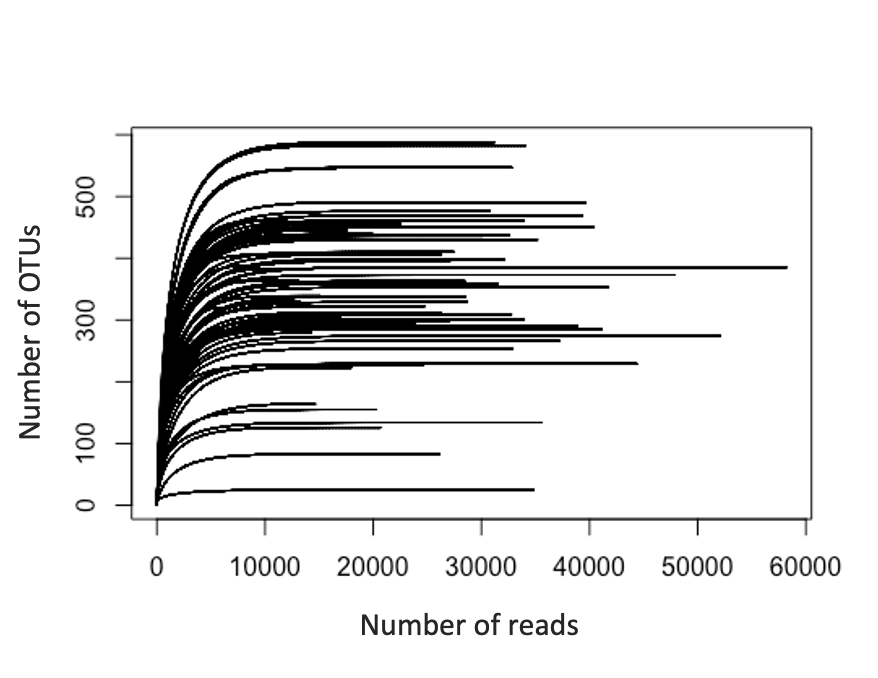

**Figure S2.** Relationship between the number of sequence reads and OTU numbers, i.e., rarefaction curves for the samples.

(Figure S3)

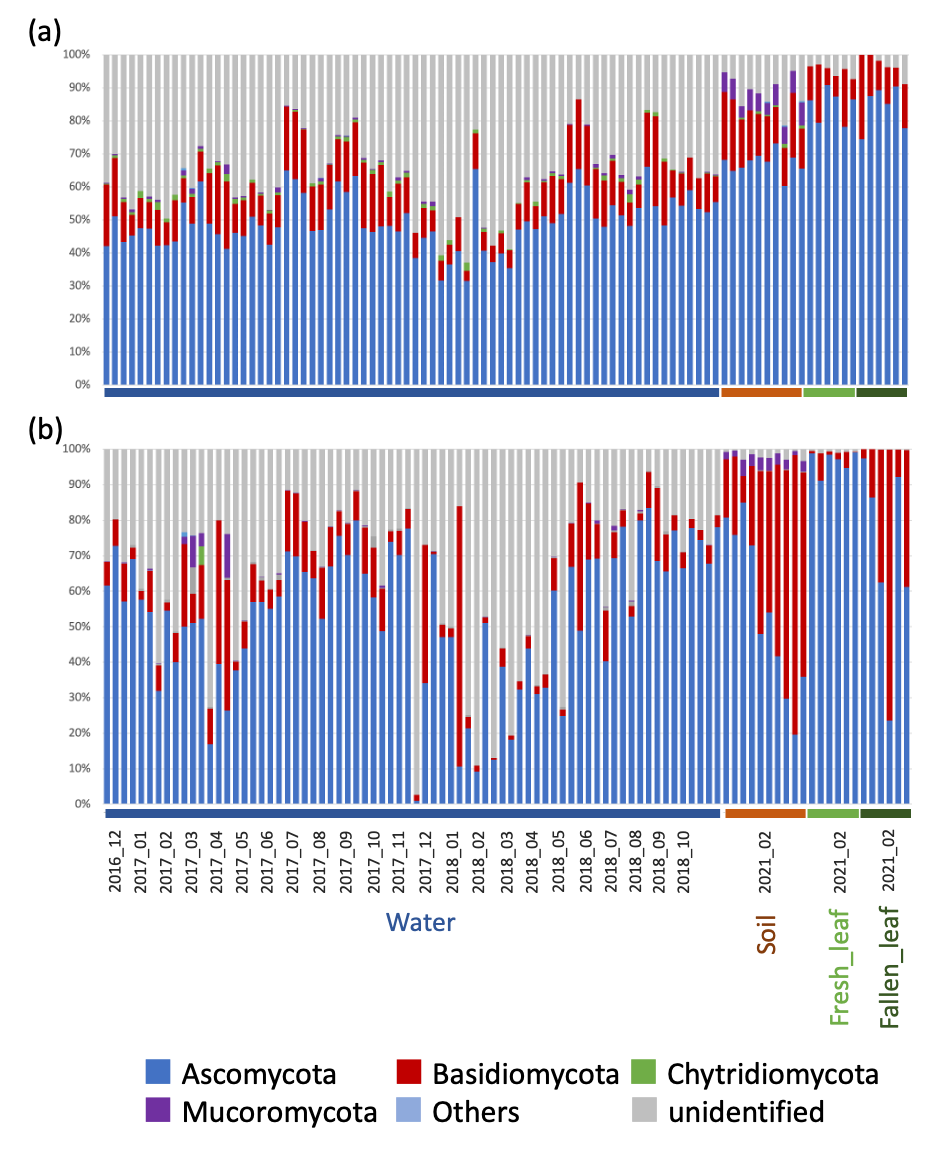

**Figure S3.** Phylum level proportions of the DNA assemblages of each sample. (a) Percentage of the number of OTUs, and (b) percentage of the number of reads in the sequence. The order of the samples is the same as that in Table S2.

(Figure S4)

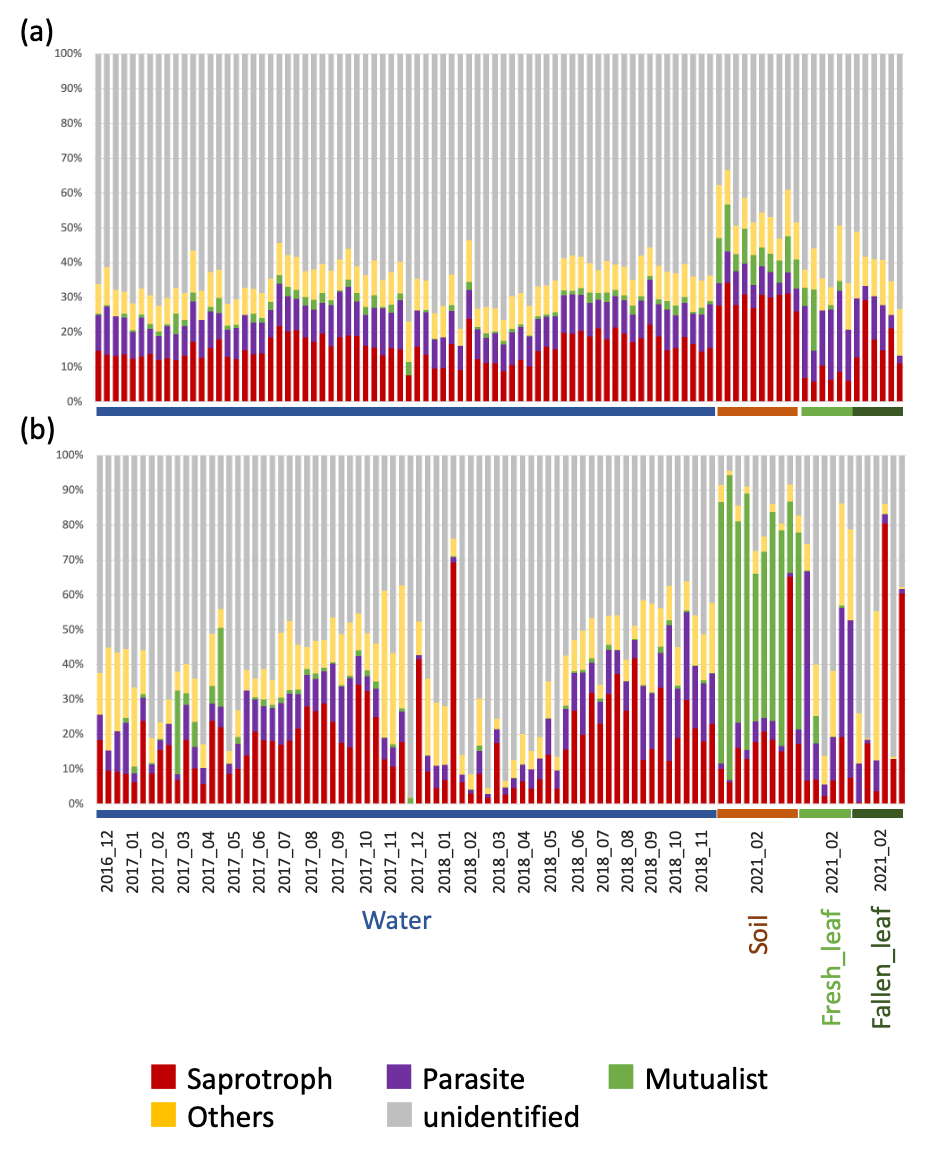

**Figure S4.** Functional composition of the DNA assemblages of each sample. (a) Percentage of the number of OTUs, and (b) percentage of the number of reads in the sequence. The order of the samples is the same as that in Table S2.

(Figure S5)

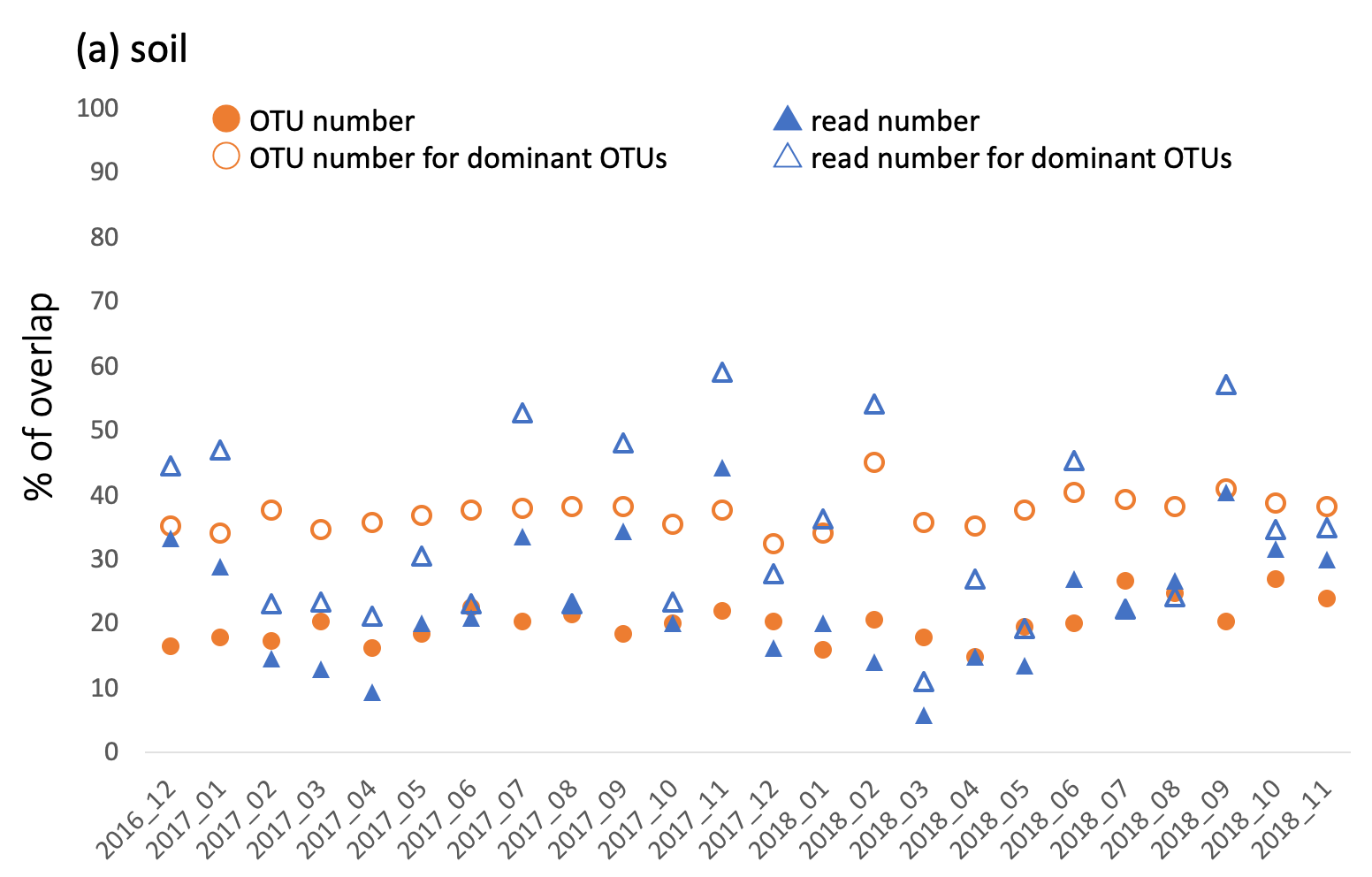

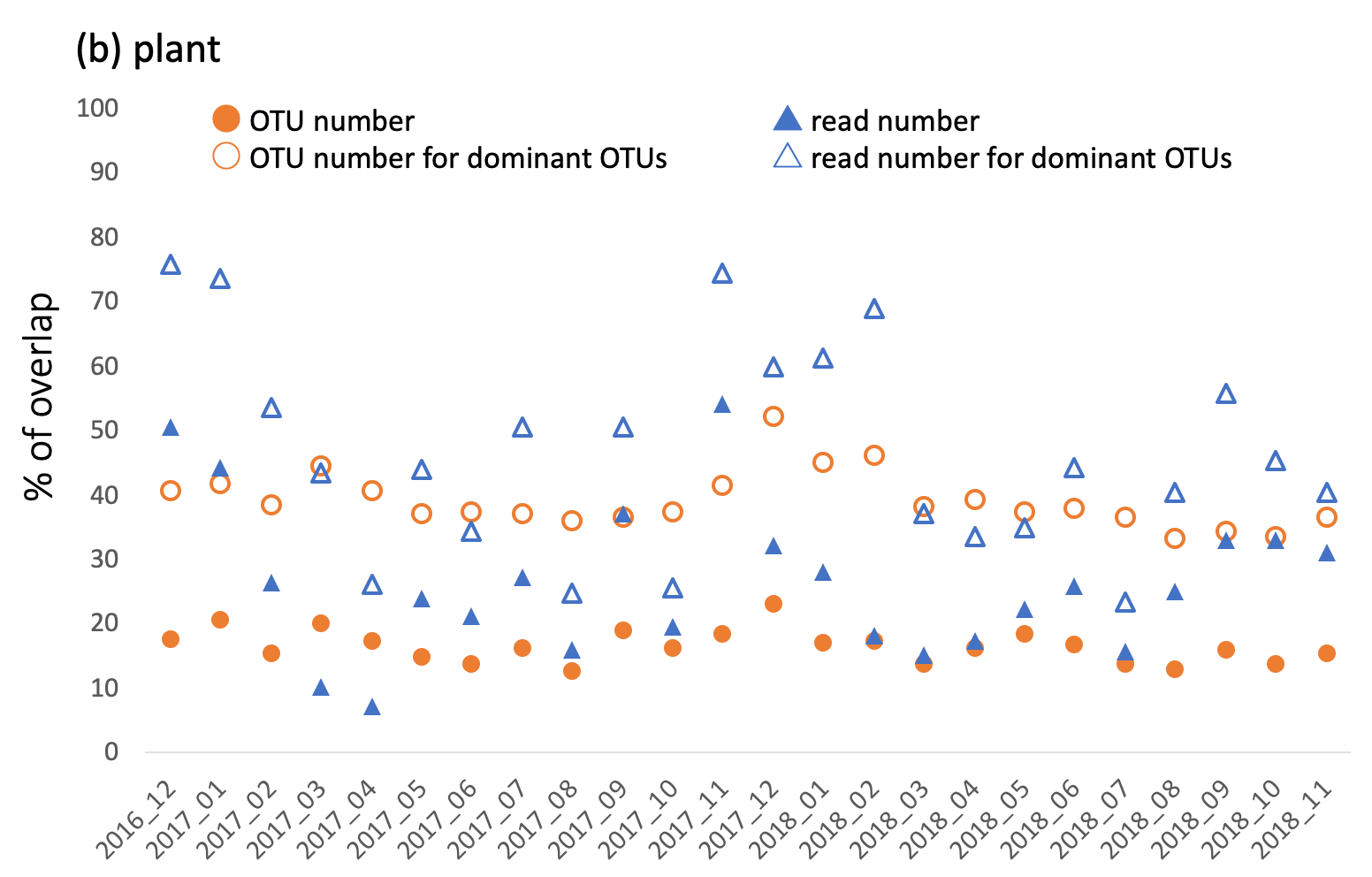

**Figure S5.** Percentage of OTUs and reads detected in stream water that overlapped with OTUs and reads detected in (a) soils and (b) plant leaves. The results are shown for the dataset of all OTUs detected in the water and the dataset of dominant OTUs (top 100 OTUs with high detection frequency).

(Figure S6)

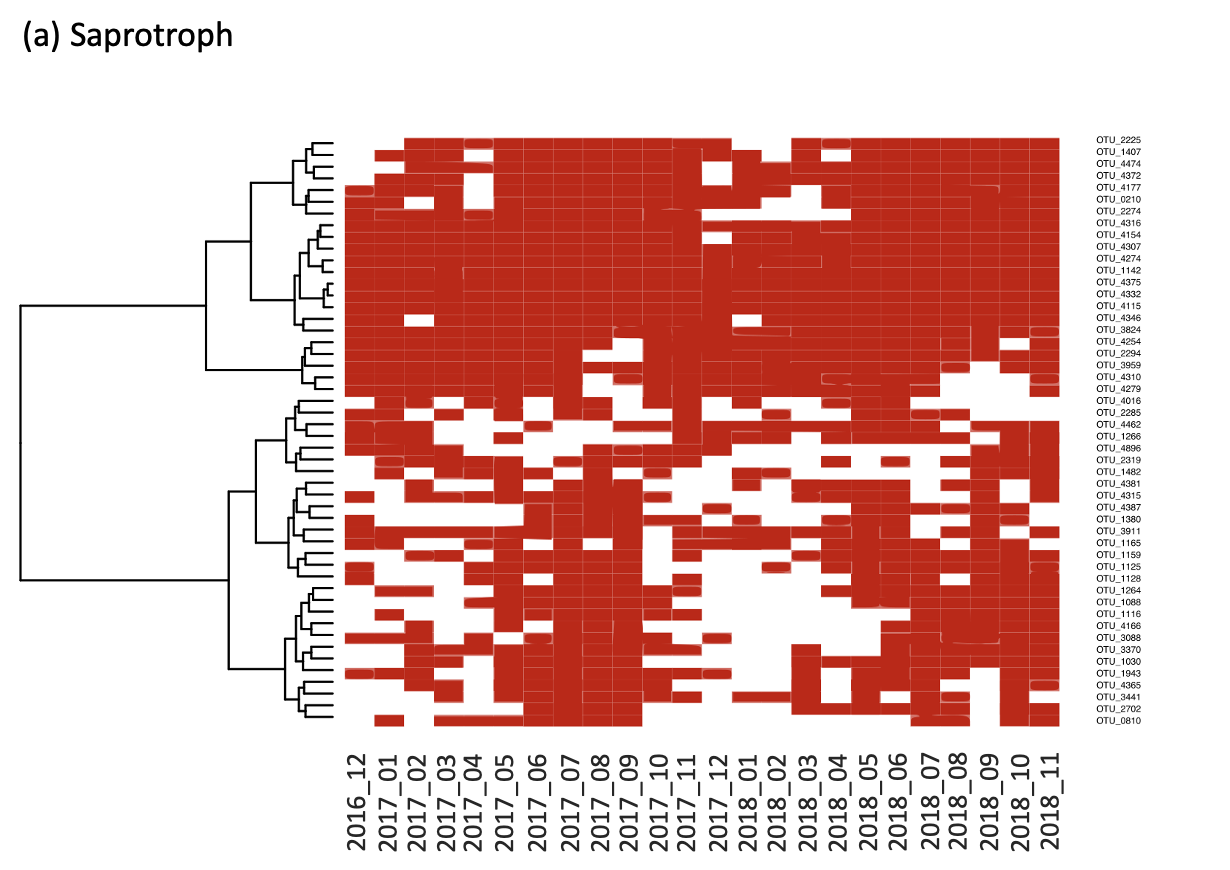

**Figure S6.** Heatmaps showing the temporal occurrence patterns of the top 50 OTUs, with the number of sample occurrences for each guild. The OTUs were clustered using the word method based on their occurrence patterns. (a) Saprotroph, (b) Parasite, and (c) Mutualist.

(Figure S6)

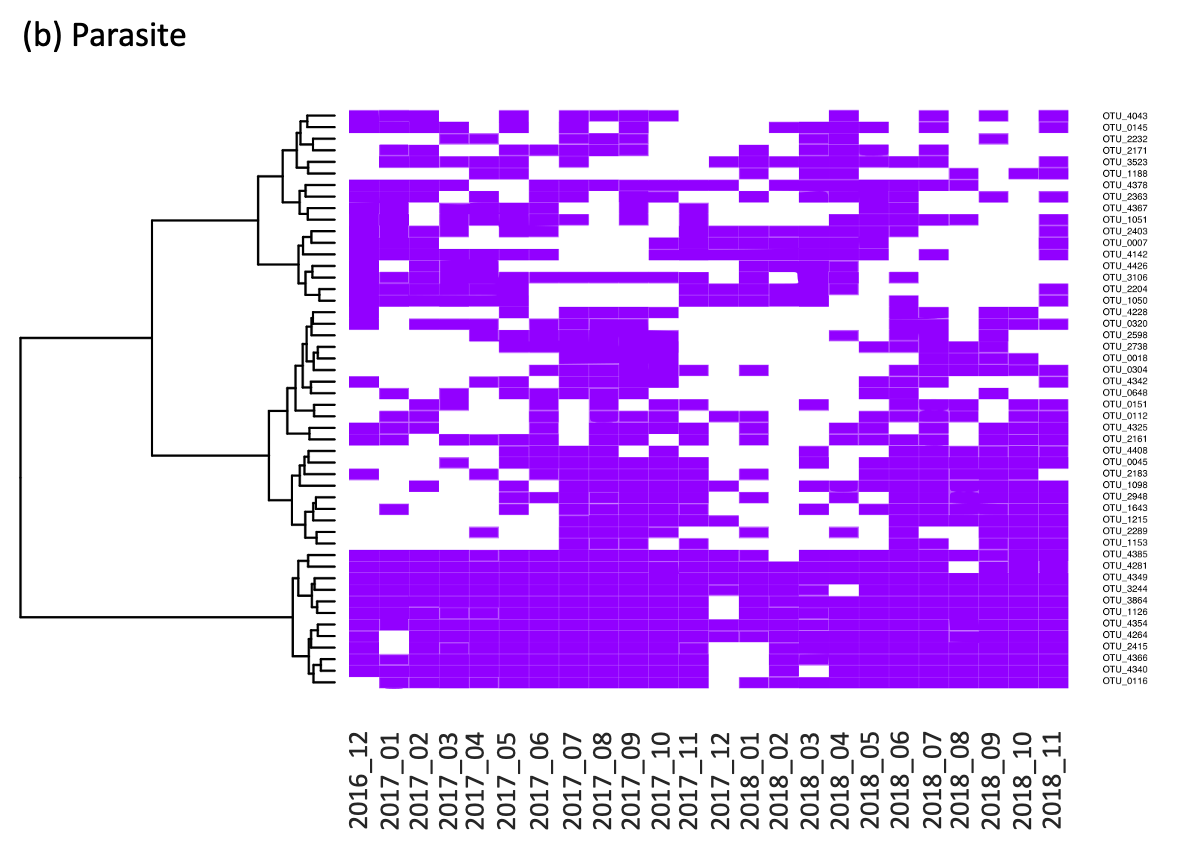

(Figure S6)

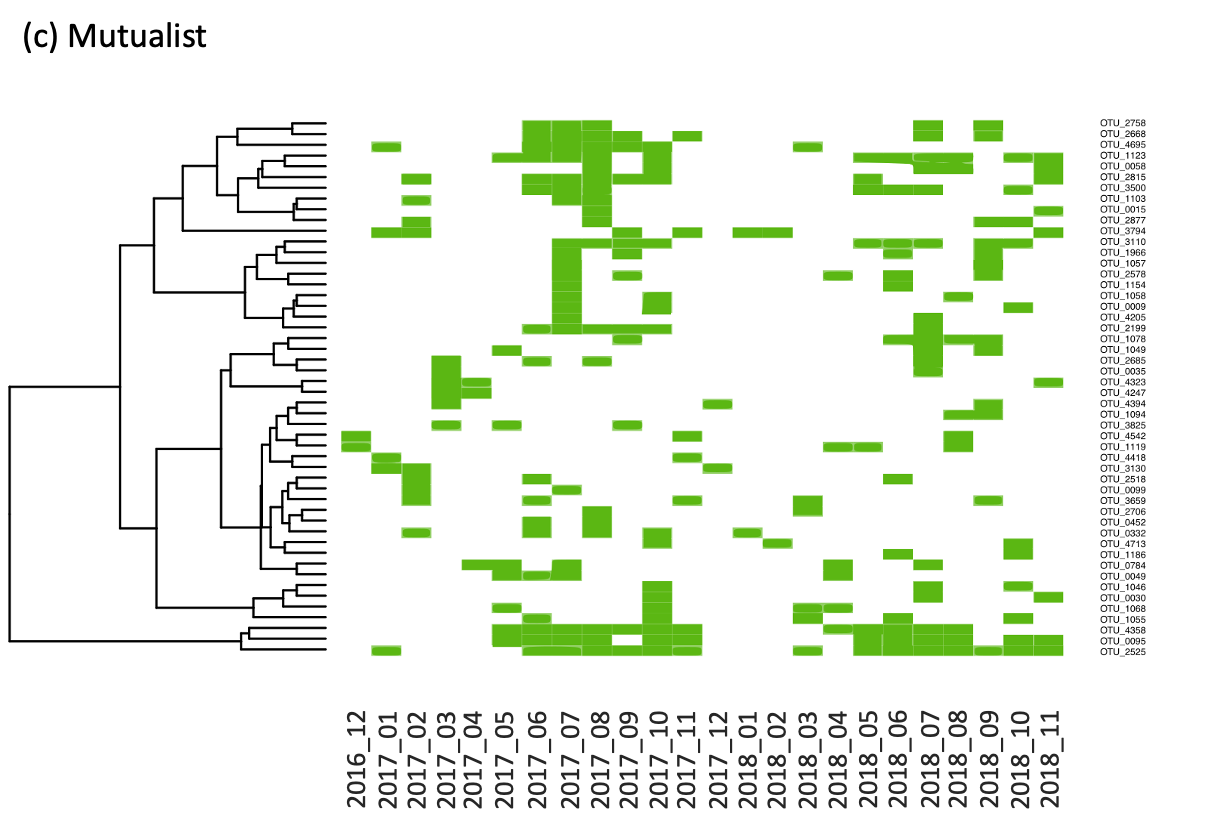

(Figure S7)

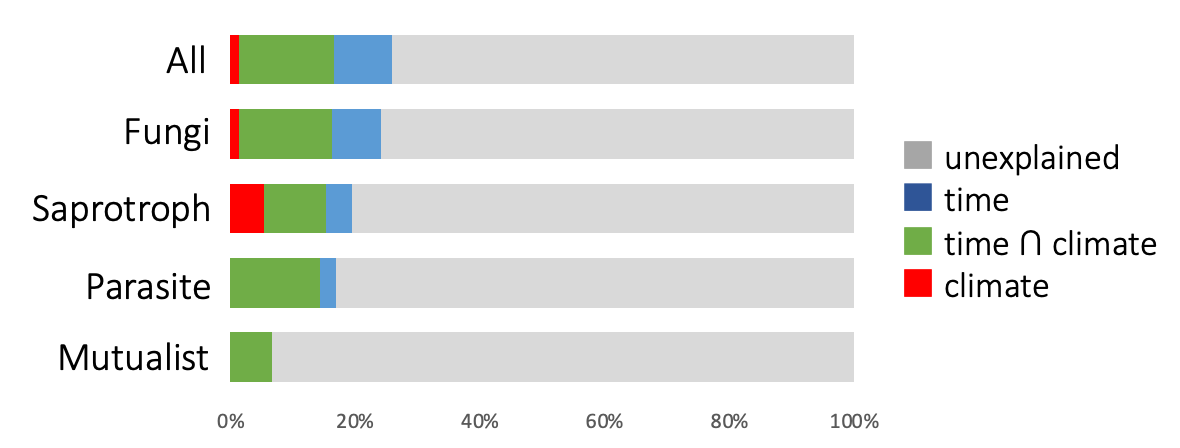

**Figure S7.** Bar plots showing pure and shared effects of climatic and temporal variables on the fungal OTU assemblages, as derived from variation partitioning analysis. Numbers indicate the proportions of explained variations. "All" shows the results for the dataset containing all detected OTUs, and "Fungi" shows the results for the dataset containing only OTUs assigned as fungi.
